## Supplementary material for "Rvb1/Rvb2 proteins couple transcription and translation during glucose starvation": Figure Supplements + Tables

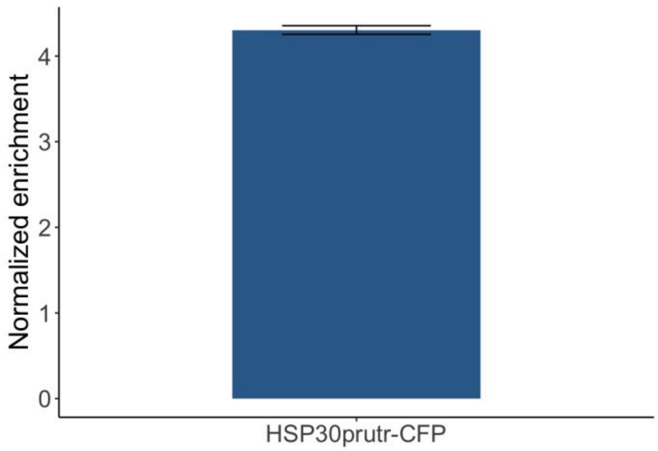


**Figure 1-figure supplement 1: Reporter RNA was enriched upon CoTrIP plasmid immunoprecipitation.** *HSP30* promoter-driven reporter mRNA was tested via RT-qPCR. Y-axis: Ct value of the reporter mRNA was first normalized by the housekeeping gene *PDC1* and further normalized by the promoter-less CoTrIP plasmid negative control.

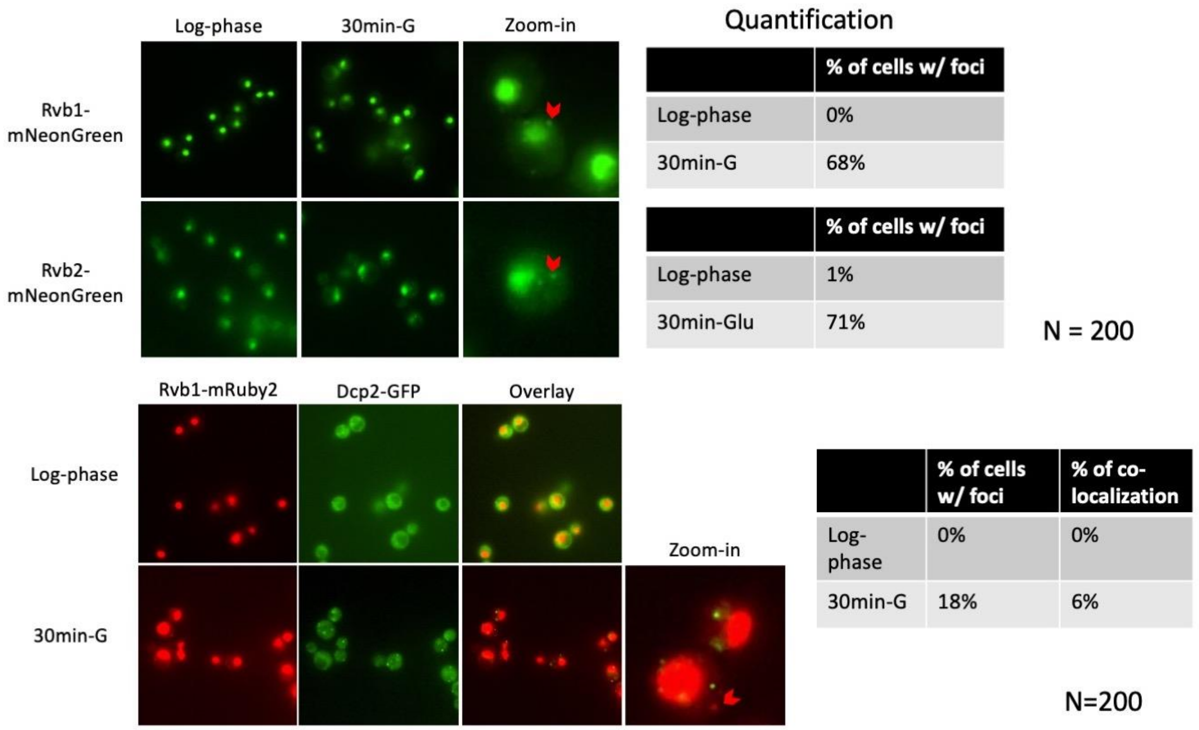


**Figure 1-figure supplement 2: Rvb1/Rvb2 form cytoplasmic granules that are not co-localized with P-body during glucose starvation.** In the upper panel, Rvb1/Rvb2 is C-terminally fused with green fluorescent protein mNeonGreen. In the lower panel, Rvb1/Rvb2 is C-terminally fused with red fluorescent protein mRuby2 and P-body marker Dcp2 is C-terminally fused with GFP. Cells are imaged in both log phase and 30-minute glucose starvation conditions. Quantification was performed on 200 cells in each imaging experiment. % of cells w/ foci: among the cells analyzed, the percentage of cells that have a Rvb-containing foci. % of co-localization: among the cells with the Rvb-containing foci, the percentage of cells that have a co-localized foci with Dcp2-containing foci.

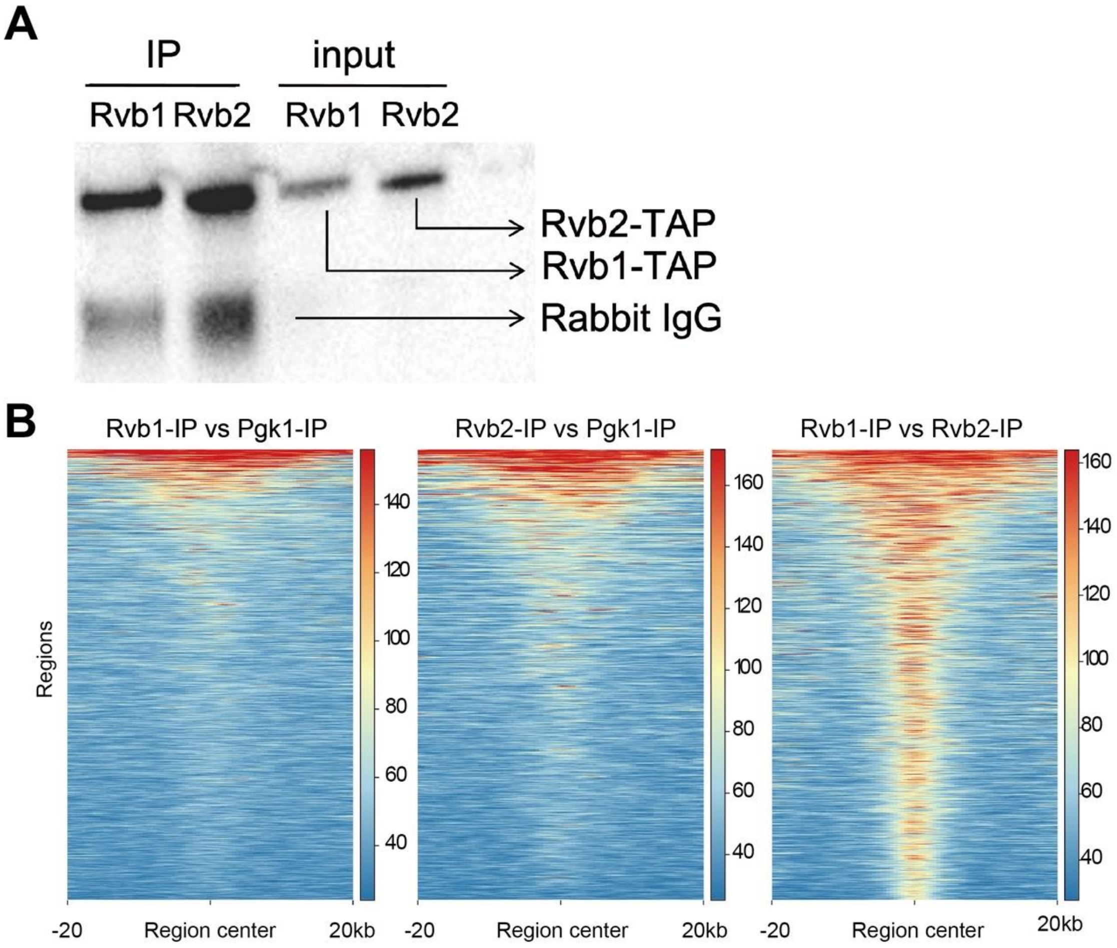


**Figure 2-figure supplement 3: Western validation of ChIP-seq, and Rvb1 and Rvb2's enrichment regions are highly overlapped.** (A) Western validating the efficiency of the immunoprecipitation (IP). Rvb1/Rvb2 are C-terminally fused with the tandem-affinity-purification tag (TAP), labeled as Rvb1-TAP and Rvb2-TAP. The proteins of interest were pulled down by rabbit IgG. Cells are harvested at 10-minute glucose starvation. Input is 1% of the total lysate. (B) comparison analysis of Rvb1/Rvb2/Pgk1's enrichment on the genome. X-axis: the enriched regions aligned by the center. Y-axis: regions arranged high to low by the level of overlapping. Color-code: red indicates that the two IPs are highly overlapped and blue means not overlapped. Left panel: use Pgk1's enriched regions as reference, score Rvb1's enrichment. Middle panel: use Pgk1's enriched regions as reference, score Rvb2's enrichment. Right panel: use Rvb2's enriched regions as reference, score Rvb1's enrichment.

#### Class I Upregulated High-Ribo genes

| Systematic Name | Name | BY4741 | | EY0690 | |
|---|---|---|---|---|---|
| | | mRNA | Ribo | mRNA | Ribo |
| YCR021C | *HSP30* | 5.9 | 2.5 | 6.9 | 0.8 |
| YBR072W | *HSP26* | 3.9 | 1.1 | 4.9 | 0.3 |
| YLL026W | *HSP104* | 3.8 | 0.7 | 3.1 | 0.3 |
| YDR258C | *HSP78* | 3.5 | 0.3 | 2.6 | -0.2 |
| YFL014W | *HSP12* | 3.1 | 0.8 | 2.6 | 0.9 |
| YPL240C | *HSP82* | 2.8 | 0.1 | 2.2 | 0.1 |
| YER067W | *RGI1* | 6.7 | 0.6 | 4.6 | -0.3 |
| YLR327C | *TMA10* | 5.9 | 2.9 | 6.0 | 0.9 |
| YKR075C | | 4.6 | 1.3 | 4.4 | -0.4 |
| YGR142W | *BTN2* | 5.0 | 2.0 | 5.2 | 0.6 |
| YBL078C | *ATG8* | 2.8 | 0.8 | 2.2 | 1.1 |

#### Class II Upregulated Low-Ribo genes

| Systematic Name | Name | BY4741 | | EY0690 | |
|---|---|---|---|---|---|
| | | mRNA | Ribo | mRNA | Ribo |
| YEL011W | *GLC3* | 5.5 | -1.8 | 5.0 | -2.7 |
| YFR015C | *GSY1* | 6.0 | -1.7 | 3.7 | -2.6 |
| YFR053C | *HXK1* | 5.5 | -1.9 | 3.5 | -1.3 |
| YPR160W | *GPH1* | 4.2 | -1.8 | 3.6 | -2.0 |
| YGR088W | *CTT1* | 3.9 | -1.0 | 4.4 | -1.2 |
| YGL205W | *POX1* | 4.1 | -1.3 | 2.8 | -1.8 |
| YDR171W | *HSP42* | 5.8 | -2.0 | 4.7 | -1.5 |
| YGR249W | *MGA1* | 5.1 | -1.8 | 5.4 | -2.2 |
| YJL052W | *TDH1* | 2.9 | -1.5 | 2.7 | -1.3 |
| YBR147W | *RTC2* | 2.5 | -2.9 | 3.0 | -2.6 |
| YKL217W | *JEN1* | 6.0 | -1.6 | 5.0 | -1.5 |
| YLR258W | *GSY2* | 3.3 | -1.1 | 1.4 | -1.1 |
| YCR091W | *KIN82* | 2.6 | -2.1 | 2.9 | -1.9 |

**Figure 2-figure supplement 4: List of Class I upregulated and high-ribo genes and Class II upregulated and low-ribo genes.** Data from Zid & O'Shea, 2014. Fold-change in mRNA levels and in ribosome occupancy after 15 minutes of glucose starvation from one measurement of BY4741 and one measurement of EY0690. mRNA: log2 mRNA fold change for glucose starvation vs. log phase glucose-rich. Ribo: log2 ribosome occupancy fold change for glucose starvation vs. log phase glucose-rich.

| Gene | FC Rvb1 vs Pgk1 | FDR% | FC Rvb2 vs Pgk1 | FDR% |
|---|---|---|---|---|
| YDR171W | 2.23 | 0.00 | 2.38 | 0.00 |
| YELOIIW | 3.43 | 0.00 | 3.58 | 0.00 |
| YFR015C | 3.57 | 0.00 | 3.07 | 0.00 |
| YFR053C | 4.13 | 0.00 | 3.45 | 0.00 |
| YGR088W | 3.34 | 0.00 | 2.85 | 0.00 |
| YGR249W | 2.81 | 0.00 | 2.75 | 0.00 |
| YJL052W | 3.5 | 0.00 | 4.26 | 0.00 |
| YPR160W | 3.97 | 0.00 | 3.52 | 0.00 |

**Figure 2-figure supplement 5: List of Rvb1/Rvb2 peak calls on the genome.** MACS algorithm was applied from the ChIP-seq results of Rvb1/Rvb2 in 10-minute glucose starvation.. Genes are showed under systematic names. FC: fold change of Rvb's peak versus Pgk1's peak. FDR: false discovery rate.

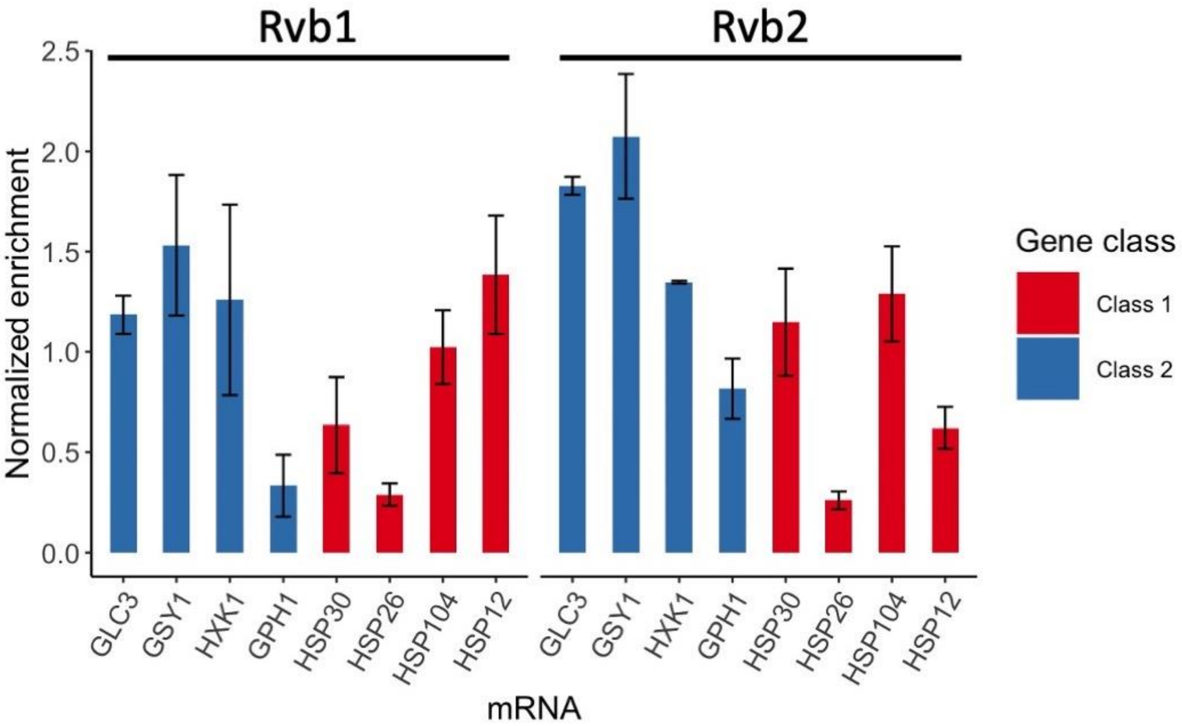


**Figure 3-figure supplement 6: Rvb1/Rvb2 did not show differential enrichment between Class I and Class II mRNAs in glucose-rich log-phase cells.** RNA immunoprecipitation qPCR of cells in log phase. Error bars are from 2 technical replicates. X-axis: 4 Class I mRNAs labeled in red and 4 Class II mRNAs in blue. Y-axis: Ct values were firstly normalized by internal control *ACT1*, then normalized by input control, finally normalized by the wild-type immunoprecipitation control group. Input: 1% of the cell lysate.

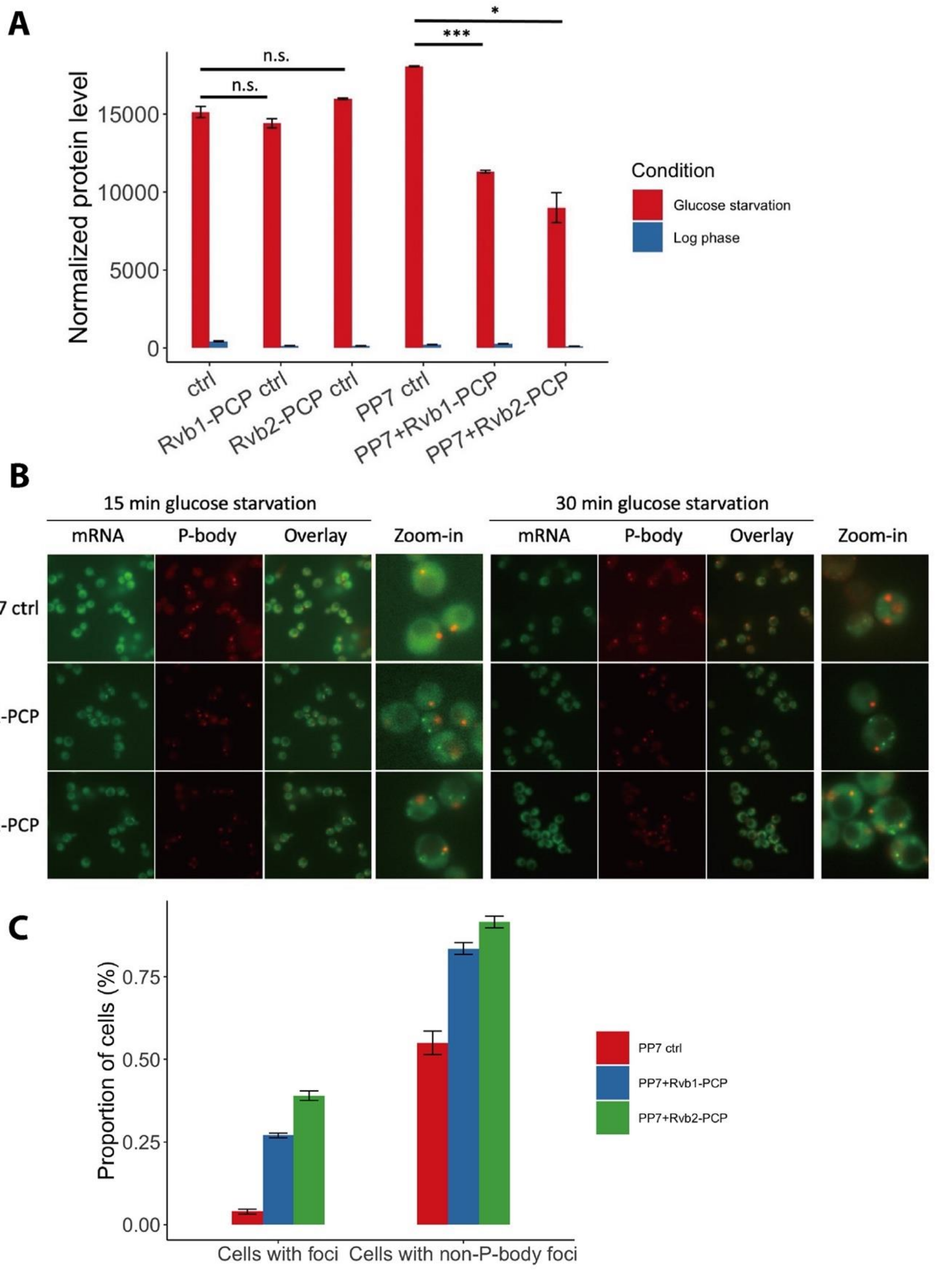
A
Normalized protein level
15000
10000
5000
0
n.s.
n.s.
***
*
ctrl
Rvb1-PCP ctrl
Rvb2-PCP ctrl
PP7 ctrl
PP7+Rvb1-PCP
PP7+Rvb2-PCP
Condition
Glucose starvation
Log phase
B
15 min glucose starvation
30 min glucose starvation
mRNA
P-body
Overlay
Zoom-in
PP7 ctrl
PP7+Rvb1-PCP
PP7+Rvb2-PCP
C
Proportion of cells (%)
0.75
0.50
0.25
0.00
Cells with foci
Cells with non-P-body foci
PP7 ctrl
PP7+Rvb1-PCP
PP7+Rvb2-PCP

**Figure 4-figure supplement 7: Engineered Rvb1/Rvb2 tethering to *HSP30* promoter-driven reporter mRNA directs cytoplasmic granular localization and repressed translation.** (A) protein synthesis of *HSP30* promoter-driven reporter mRNA in log phase and 25-minute glucose starvation. Y-axis: nanoluciferase synthesized within 5-minute time frame. Nanoluciferase reading was subtracted by the nanoluciferase reading of cycloheximide added 5 minutes earlier. X-axis: different Rvb-tethering conditions. Ctrl: negative control, no PP7 and PP7-coat protein. Rvb1-PCP ctrl: negative control, Rvb1 is fused with PP7-coat protein but no reporter mRNA with PP7 loop. Rvb2-PCP ctrl: negative control, Rvb2 is fused with PP7-coat protein but no reporter mRNA with PP7 loop. PP7 ctrl: negative control, cells only have the reporter mRNA with PP7 loop. PP7+Rvb1-PCP: Rvb1 is tethered to mRNA. PP7+Rvb2-PCP: Rvb2 is tethered to mRNA. Log phase is labeled in blue and glucose starvation in red. Error bars are from 2 biological replicates. Statistical significance was assessed by 2-sample t-test. Null hypothesis: experimental groups and control groups have equivalent results (*$p<0.05$, **$p<0.01$, ***$p<0.001$). (B) Live imaging showing the subcellular localization of the *HSP30* promoter-driven reporter mRNA in 15-minute and 30-minute glucose starvation. Reporter mRNA is labeled by the MS2 imaging system. P-body is labeled by marker protein Dcp2. PP7 ctrl: negative control, cells only have the reporter mRNA with PP7 loop. PP7+Rvb1-PCP: Rvb1 is tethered to mRNA. PP7+Rvb2-CP: Rvb2 is tethered to mRNA. (C) Quantification of the subcellular localization of the reporter mRNA in 30-minute glucose starvation. Cells with foci: percentage of cells that have the reporter mRNA-containing granule foci. Cells with non-P-body foci: among the cells that have the reporter mRNA-containing granule foci, the percentage of cells that have reporter mRNA-containing granule foci that are not co-localized with P-body. N=200. Error bars are from 2 biological replicates.

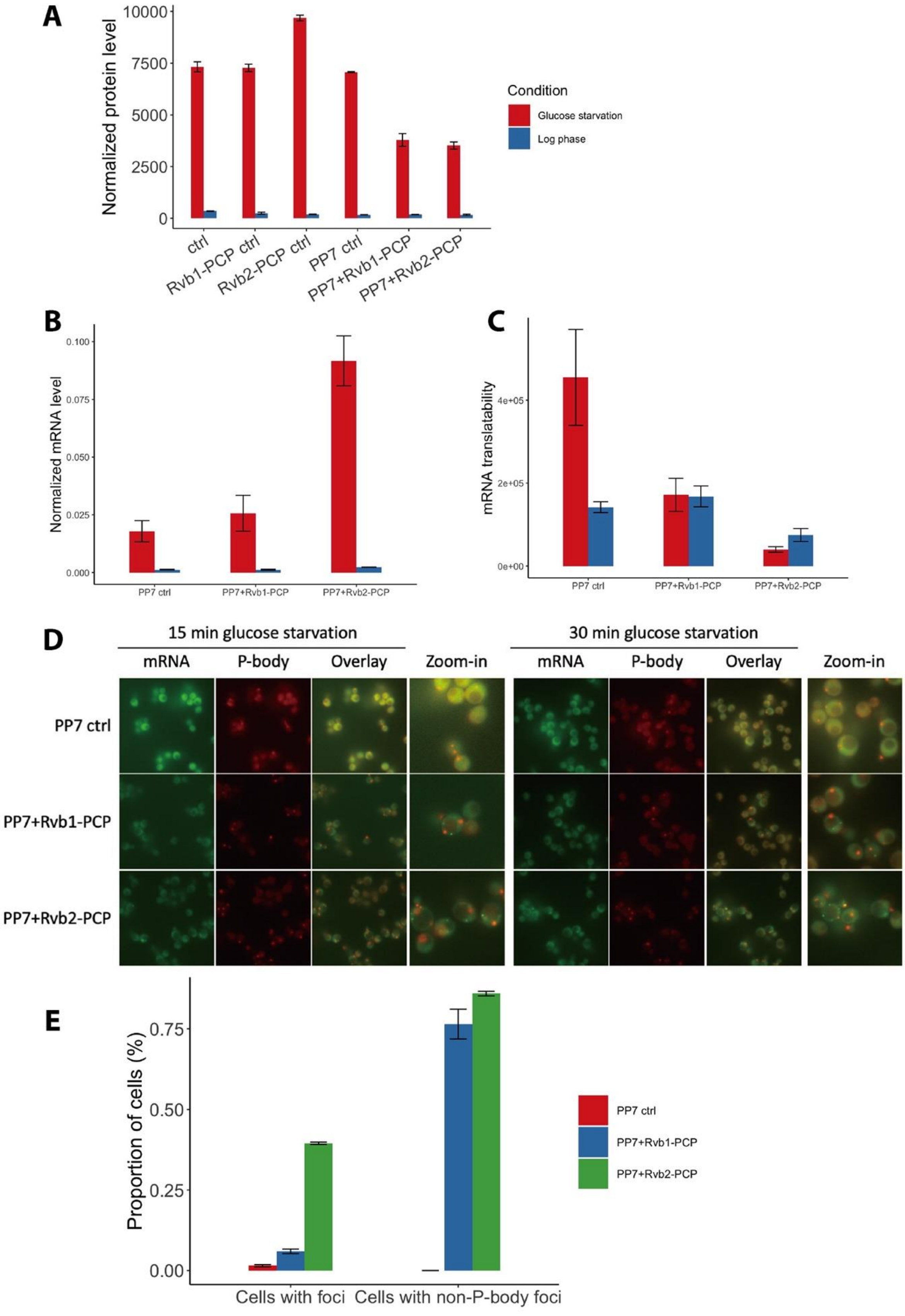
A
Normalized protein level
10000
7500
5000
2500
0
ctrl
Rvb1-PCP ctrl
Rvb2-PCP ctrl
PP7 ctrl
PP7+Rvb1-PCP
PP7+Rvb2-PCP
Condition
Glucose starvation
Log phase
B
Normalized mRNA level
0.100
0.075
0.050
0.025
0.000
PP7 ctrl
PP7+Rvb1-PCP
PP7+Rvb2-PCP
C
mRNA translatability
4e+05
2e+05
0e+00
PP7 ctrl
PP7+Rvb1-PCP
PP7+Rvb2-PCP
D
15 min glucose starvation
30 min glucose starvation
mRNA
P-body
Overlay
Zoom-in
PP7 ctrl
PP7+Rvb1-PCP
PP7+Rvb2-PCP
E
Proportion of cells (%)
0.75
0.50
0.25
0.00
Cells with foci
Cells with non-P-body foci
PP7 ctrl
PP7+Rvb1-PCP
PP7+Rvb2-PCP

**Figure 4-figure supplement 8: Engineered Rvb1/Rvb2 tethering to *HSP26* promoter-driven reporter mRNA directs cytoplasmic granular localization and repressed translation.** (A) protein synthesis of *HSP26* promoter-driven reporter mRNA in log phase and 25-minute glucose starvation. Y-axis: nanoluciferase synthesized within 5-minute time frame. Nanoluciferase reading was subtracted by the nanoluciferase reading of cycloheximide added 5 minutes earlier. (B) mRNA levels of *HSP26* promoter-driven reporter mRNA in log phase and 15-minute glucose starvation. Y-axis: Ct values of reporter mRNAs were normalized by the internal control *ACT1*. (C) Translatability of *HSP26* promoter-driven reporter mRNA in log phase and 15-minute glucose starvation. mRNA translatability: normalized protein level over normalized mRNA level. (A, B, C) Log phase is labeled in blue and glucose starvation in red. Error bars are from 2 biological replicates. Ctrl: negative control, no PP7 and PP7-coat protein. Rvb1-PCP ctrl: negative control, Rvb1 is fused with PP7-coat protein but no reporter mRNA with PP7 loop. Rvb2-PCP ctrl: negative control, Rvb2 is fused with PP7-coat protein but no reporter mRNA with PP7 loop. PP7 ctrl: negative control, cells only have the reporter mRNA with PP7 loop. PP7+Rvb1-PCP: Rvb1 is tethered to mRNA. PP7+Rvb2-PCP: Rvb2 is tethered to mRNA. (D) Live imaging showing the subcellular localization of the *HSP26* promoter-driven reporter mRNA in 15-minute and 30-minute glucose starvation. Reporter mRNA is labeled by the MS2 imaging system. P-body is labeled by marker protein Dcp2. PP7 ctrl: negative control, cells only have the reporter mRNA with PP7 loop. PP7+Rvb1-PCP: Rvb1 is tethered to mRNA. PP7+Rvb2-CP: Rvb2 is tethered to mRNA. (E) Quantification of the subcellular localization of the reporter mRNA in 30-minute glucose starvation. Cells with foci: percentage of cells that have the reporter mRNA-containing granule foci. Cells with non-P-body foci: among the cells that have the reporter mRNA-containing granule foci, the percentage of cells that have reporter mRNA-containing granule foci that are not co-localized with P-body. N=200. Error bars are from 2 biological replicates.

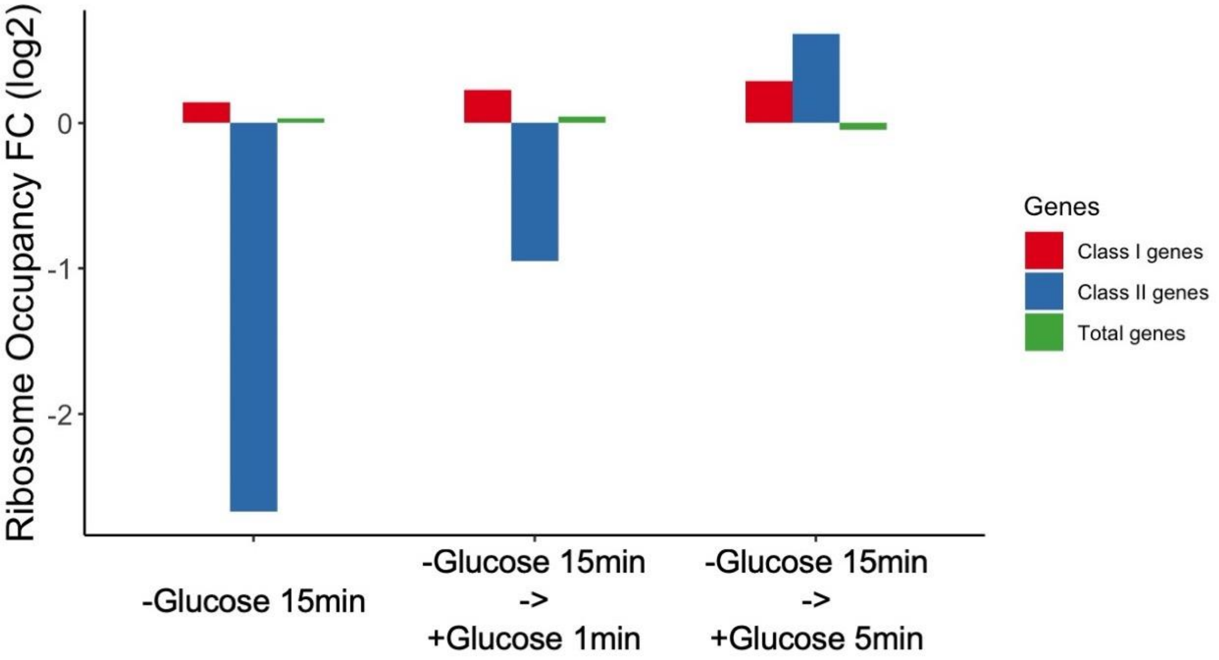


**Figure 4-figure supplement 9: Ribosome occupancy of endogenous glucose metabolism mRNAs was quickly induced after glucose replenishment.** Ribosome profiling was performed on cells in 15-minute glucose starvation and followed by 1-minute and 5-minute glucose addback. Y-axis: log 2 scale of ribosome occupancy fold changes on mRNAs compared to log phase condition. X-axis: each bar represents a gene group. Bars are grouped by conditions. Class I genes and Class II genes refer to Figure 2-figure supplement 4.

### Supplementary File 1 – Strains and Plasmids

| Strain ID | Genotype | Source |
|---|---|---|
| EY0690/W303 | MATa trp1 leu2 ura3 his3 can1 GAL+ psi+ | Lab stock |
| ZY1 | EY0690; pRS406-CMV lacIA-FLAG | This study |
| ZY2 | EY0690; HSP30-CFP-CoTrIP | This study |
| ZY3 | ZY1; HSP30prUTR-CFP-CoTrIP | This study |
| ZY4 | ZY1; HXK1prUTR-CFP-CoTrIP | This study |
| ZY5 | ZY1; HSP26prUTR-CFP-CoTrIP | This study |
| ZY6 | ZY1; GLC3prUTR-CFP-CoTrIP | This study |
| ZY7 | ZY1; Blank-CoTrIP | This study |
| ZY18 | EY0690; MYO2pr-MS2-CP-GFP2x; Dcp2-RFP | Zid & O'Shea 2014 |
| ZY147 | EY0690; Rvb1-TAP | This study |
| ZY148 | EY0690; Rvb2-TAP | This study |
| ZY282 | EY0690; Dcp2-GFP; Rvb1-mRuby2 | This study |
| ZY284 | EY0690; Dcp2-GFP; Rvb2-mRuby2 | This study |
| ZY266 | ZY18; HSP30prUTR-nLuc-pest-1XPP7-12XMS2-tADH1 | This study |
| ZY314 | ZY18; Rvb1-PP7CP-6Xhis; HSP30prUTR-nLuc-pest-1XPP7-12XMS2-tADH1 | This study |
| ZY315 | ZY18; Rvb2-PP7CP-6Xhis; HSP30prUTR-nLuc-pest-1XPP7-12XMS2-tADH1 | This study |
| ZY193 | ZY18; HSP30prUTR-nLuc-pest-12XMS2-tADH1 | This study |
| ZY446 | ZY18; Rvb1-PP7CP-6Xhis; HSP30prUTR-nLuc-pest-12XMS2-tADH1 | This study |
| ZY449 | ZY18; Rvb2-PP7CP-6Xhis; HSP30prUTR-nLuc-pest-12XMS2-tADH1 | This study |
| ZY316 | ZY18; Rvb1-PP7CP-6Xhis | This study |
| ZY317 | ZY18; Rvb2-PP7CP-6Xhis | This study |
| ZY318 | EY0690; Rvb1-mNeogreen | This study |
| ZY319 | EY0690; Rvb2-mNeogreen | This study |
| ZY488 | ZY18; HSP26prUTR-nLuc-pest-12XMS2-tADH1 | This study |
| ZY489 | ZY18; HSP12prUTR-nLuc-pest-12XMS2-tADH1 | This study |
| ZY490 | ZY18; HSP26prUTR-nLuc-pest-1XPP7-12XMS2-tADH1 | This study |
| ZY491 | ZY18; HSP12prUTR-nLuc-pest-1XPP7-12XMS2-tADH1 | This study |
| ZY492 | ZY18; HSP26prUTR-nLuc-pest-12XMS2-tADH1; Rvb1-PP7CP-6Xhis | This study |
| ZY493 | ZY18; HSP12prUTR-nLuc-pest-12XMS2-tADH1; Rvb1-PP7CP-6Xhis | This study |
| ZY494 | ZY18; HSP26prUTR-nLuc-pest-1XPP7-12XMS2-tADH1; Rvb1-PP7CP-6Xhis | This study |
| ZY495 | ZY18; HSP12prUTR-nLuc-pest-1XPP7-12XMS2-tADH1; Rvb1-PP7CP-6Xhis | This study |
| ZY496 | ZY18; HSP26prUTR-nLuc-pest-12XMS2-tADH1; Rvb2-PP7CP-6Xhis | This study |
| ZY497 | ZY18; HSP12prUTR-nLuc-pest-12XMS2-tADH1; Rvb2-PP7CP-6Xhis | This study |

| | | |
|---|---|---|
| ZY498 | ZY18; HSP26prUTR-nLuc-pest-1XPP7-12XMS2-tADH1; Rvb2-PP7CP-6Xhis | This study |
| ZY499 | ZY18; HSP12prUTR-nLuc-pest-1XPP7-12XMS2-tADH1; Rvb2-PP7CP-6Xhis | This study |
| ZY642 | EY0690; HSP26prUTR-CFP-12XMS2-tADH1; Rvb1-TAP | This study |
| ZY643 | EY0690; GLC3prHSP26UTR-CFP-12XMS2-tADH1; Rvb1-TAP | This study |
| ZY644 | EY0690; HSP26prUTR-CFP-12XMS2-tADH1; Rvb2-TAP | This study |
| ZY645 | EY0690; GLC3prHSP26UTR-CFP-12XMS2-tADH1; Rvb2-TAP | This study |

| Plasmid ID | Annotation | Source |
|---|---|---|
| ZP66 | pUC-TalO8 (Blank-CoTrIP) | This study |
| ZP67 | TalO8-HSP30-CFP | This study |
| ZP68 | TalO8-HXK1-CFP | This study |
| ZP69 | TalO8-HSP26-CFP | This study |
| ZP70 | TalO8-GLC3-CFP | This study |
| ZP64 | pRS406-CMV-LacI-3xFLAG | Unnikrishnan et al 2012 |
| ZP60 | pFA6-TAP(CBP-TEV-ZZ)-Kan | lab stock |
| ZP61 | pFA6-TAP(CBP-TEV-ZZ)-His | lab stock |
| ZP47 | pKT-mNeongreen-Ura | lab stock |
| ZP224 | pFA6a-link-yoEGFP-SpHis5 | lab stock |
| ZP109 | pKT-mRuby2-HPH | lab stock |
| ZP296 | pRS305-HSP30prUTR-nLuc-pest-1XPP7-12XMS2-tADH1 | This study |
| ZY311 | pKT-PP7CP-6xHis-tADH1 | lab stock |
| ZP207 | pRS305-HSP30prUTR-nLuc-PEST-12XMS2-tADH1 | This study |
| ZP214 | pRS305-GLC3prUTR-nLuc-PEST-12XMS2-tADH1 | This study |
| ZP315 | pRS305-GSY1prUTR-nLuc-PEST-12XMS2-tADH1 | This study |
| ZP441 | pRS305-HSP26prUTR-nLuc-pest-12XMS2-tADH1 | This study |
| ZP442 | pRS305-HSP12prUTR-nLuc-pest-12XMS2-tADH1 | This study |
| ZP443 | pRS305-HSP26prUTR-nLuc-pest-1XPP7-12XMS2-tADH1 | This study |
| ZP444 | pRS305-HSP12prUTR-nLuc-pest-1XPP7-12XMS2-tADH1 | This study |
| ZP440 | pRS305-1XPP7-12XMS2-tADH1 | lab stock |
| ZP29 | pRS305-HSP26prUTR-CFP-12XMS2-tADH1 | Zid & O'Shea, 2014 |
| ZP32 | pRS305-GLC3prHSP26UTR-CFP-12XMS2-tADH1 | Zid & O'Shea, 2014 |
| ZP15 | pRS305-12xMS2-tADH1 | Zid & O'Shea, 2014 |

**Supplementary File 2 – Oligos**

Cloning Oligos

| Oligo name | Description | Sequence |
|---|---|---|
| ZO464_NLuc+PestR | amplify nLuc-pest to assemble into reporter vector | ATCCACTAGTTCTAGAGC TTAAACATTAATACGAGCAGAAG |
| ZO463_yNLucF | amplify nLuc-pest to assemble into reporter vector | ATGGTTTTTACTTTAGAAGATTTTG |
| ZO966_HSP30pr-F | amplify HSP30 promoter and UTR to assemble into reporter vector | TCACTATAGGGCGAATTGGAGCTCCACCGC CCTTTCTTCAAAAGTAGAAAACTTG |
| ZO470_HSP30utr-R | amplify HSP30 promoter and UTR to assemble into reporter vector | TCTAAAGTAAAAACCAT TTGAAATTTGTTGTTTTTAGTAATCAA |
| ZO432_cRvb2-R | checking the C-terminal fusion of Rvb2 | CACCAACCAAGGCTTTTTGT |
| ZO431_cRvb2-F | checking the C-terminal fusion of Rvb2 | TGACCAAAACAGGTGTGGAA |
| ZO430_cRvb1-R | checking the C-terminal fusion of Rvb1 | CACAGCCATTACCACACCAG |
| ZO429_cRvb1-F | checking the C-terminal fusion of Rvb1 | CCTGAAGACGCAGAGAATCC |
| ZO247RVB1TAPtag_F | C-terminal TAP-tag | AAGGTCAACAAAGATTTTAGAAACTTCCGCAAATTATTTG cggatccccgggttaattaa |
| ZO246RVB1Taptag_R | C-terminal TAP-tag | TATTTTTATTTATGAAATGTGCTTTAGGCTTTCTTCACTG gaattcgagctcgtttaaac |
| ZO245RVB2TAPtag_F | C-terminal TAP-tag | TGCTAAATCAGCAGACCCTGATGCCATGGATACTACGGAAcggatccccgggttaattaa |
| ZO244RVB2TAPtag_R | C-terminal TAP-tag | TATATATTTGATGCAATTTCTGCCTTAAAGTACAAAATGCgaattcgagctcgtttaaac |
| ZO805_pKT_Rvb2_R | C-terminal tagging of pKT vector | TATATATTTGATGCAATTTCTGCCTTAAAGTACAAAATGC tcgatgaattcgagctcg |
| ZO804_pKT_Rvb2_F | C-terminal tagging of pKT vector | TGCTAAATCAGCAGACCCTGATGCCATGGATACTACGGAA ggtgacggtgctggttta |
| ZO803_pKT_Rvb1_R | C-terminal tagging of pKT vector | TATTTTTATTTATGAAATGTGCTTTAGGCTTTCTTCACTG tcgatgaattcgagctcg |
| ZO802_pKT_Rvb1_F | C-terminal tagging of pKT vector | AAGGTCAACAAAGATTTTAGAAACTTCCGCAAATTATTTG ggtgacggtgctggttta |
| ZO680_PP7_RE2 | PP7 stem loop with NotI/BamHI overhangs | GATCC TAAGGGTTTCCATATAAACTCCTTAAGC |
| ZO679_PP7_RE1 | PP7 stem loop with NotI/BamHI overhangs | GGCCGC TTAAGGAGTTTATATGGAAACCCTTA G |

qPCR Oligos

| Oligo name | Sequence |
|---|---|
| ZO807_qMS2-CP-R | GTCGGAATTCGTAGCGAAAA |
| ZO806_qMS2-CP-F | GCAGAATCGCAAATACACCA |
| ZO554_qnLuc_R | CCTTCATAAGGACGACCAAA |
| ZO553_qnLuc_F | TGGTGATCAAATGGGTCAAA |
| ZO517_qHsp12prR | GAGCGGGTAACAGATGGAAG |
| ZO516_qHsp12prF | GCGCTGCAAGTTCCTTACTT |
| ZO515_qHsp104prR | ATGAAACTCTCGCCACAACC |

| | |
|---|---|
| ZO514_qHsp104prF | AAATGGACTGGATCGACGAC |
| ZO513_qHsp26R | ATCATAAAGAGCGCCAGCAT |
| ZO512_qHsp26F | AACAGATTGCTGGGTGAAGG |
| ZO505_qGsy1prR | GCGGGAAGAAAAGAAGGAGT |
| ZO504_qGsy1prF | AGGGCAGACAAGAGGCTGTA |
| ZO84_qActR | CGGTGATTTCCTTTTGCATT |
| ZO83_qActF | CTGCCGGTATTGACCAAACT |
| qTub1prR | CGCTAGATGCATTAAACATGAAG |
| qTub1prF | GTGCTCACACCAAGCATCAT |
| qAct1prR | GAGAGGCGAGTTTGGTTTCA |
| qAct1prF | TCACCCGGCCTCTATTTTC |
| qGPH1prR | TCGTCGGTGTTCCTTCCTTA |
| qGPH1prF | GAACGCCTTCCCCAATTAC |
| qHxk1prR | CCTGGTTGCTCCAGTAAGG |
| qHxk1prF | TTCAGGAAGAATGGCAGTCC |
| qGlc3prR | TTGCAACAGCCCCTTGGAC |
| qGlc3prF | GGGCACTCATCAACAATGTG |
| qHsp26prF | CTGTCAAGGTGCATTGTTGG |
| qHsp30prR | CGGGATATGGCTTTGCTTAC |
| qHsp30prF | CGATTTTGTTGGCCATTTTCCA |
| qGsy1R | GCAGTGATTTGCGACACAGT |
| qGsy1F | GCCGCTGGTGATGTAGATTT |
| qHsp12R | TTGGTTGGGTCTTCTTCACC |
| qHsp12F | CGAAAAAGGCAAGGATAACG |
| qHsp104R | CACTTGGTTCAGCGACTTCA |
| qHsp104F | CGACGCTGCTAACATCTTGA |
| qGph1R | TCATAAGCAGCCATGTCATCA |
| qGph1F | TTCCCCAAGAAATCAAGTCAA |
| qTub1R | GGTGTAATGGCCTCTTGCAT |
| qTub1F | CCACGTTTTTCCATGAAACC |
| qHsp30R | TCAGCTTGAACACCAGTCCA |
| qHsp30F | GGGCAGTGTTTGCAGTCTTT |
| qGlc3R | CGAAATCGCCGTTAGGTAAA |
| qGlc3F | CAATCCGGAAACCAAAGAAA |
| qHsp26prF | CATAAGGGGGAGGGAATAAC |
| qCITCFPf | CTGGTGAAGGTGAAGGTGAC |
| qCITCFPr | TGTGGTCTGGGTATCTAGCG |
| qHXK1F | TTTGTAGCAATGGGACGACA |
| qHXK1R | GTACCCAGCTTCCCAAAACA |
